# Genome Mining of *Streptomyces* sp. PB17 from Cuatro Ciénegas Basin Reveals Biosynthetic Potential for Novel Neurotherapeutics

**DOI:** 10.1101/2025.02.12.637946

**Authors:** Martínez-Olivas Martha Adriana, Melchor-Rangel Silverio Alejandro, Mendoza-Cibrian Miguel Ángel, Benavides-García Paola, De la Torre-Zavala Susana

## Abstract

The Cuatro Ciénegas Basin (CCB), a biodiversity hotspot in Coahuila, Mexico, harbors microbial communities adapted to extreme oligotrophic conditions, making it a promising site for novel bioactive compound discovery.. This study reports the isolation, whole-genome sequencing, and analysis of *Streptomyces* sp. PB17, a thermotolerant actinomycete retrieved from the karstic Poza La Becerra. Phylogenomic analyses including TYGS and VBCG pipelines, combined with ANI values, support its classification as a putatively novel species. *Streptomyces* PB17’s genome mining revealed 31 biosynthetic gene clusters (BGCs), with over 10 exhibiting low similarity to known pathways, highlighting its potential for novel bioactive compounds. Among these, ribosomally synthesized and post-translationally modified peptides (RiPPs) demonstrated strong predictive scores for blood-brain barrier penetration properties, representing promising candidates for neurotherapeutic applications. These findings underscore the biotechnological potential of actinomycetes retrieved from extreme sites and emphasize the importance of underexplored ecosystems like the CCB in natural product discovery, including neurotherapeutic research.

## Introduction

The Cuatro Ciénegas Basin (CCB) in Coahuila, Mexico, is an ecological and evolutionary hotspot characterized by its oligotrophic conditions and extreme environmental pressures. These unique selective forces have driven the adaptation of diverse and largely underexplored microbial communities, (Souza *et al*. 2018) making CCB a promising site for the discovery of novel bioactive compounds (Arocha-Garza et al. 2017). Characterized by extreme environmental conditions and nutrient imbalances, such as phosphorus oligotrophy in many of its aquatic systems and soils (Elser et al. 2006), CCB offers a rare opportunity to explore life’s adaptations to resource-scarce environments (Tapia-Torres et al. 2016). Among its numerous springs and ponds, Poza La Becerra is notable for its warm waters and karstic influx from the surrounding mountains (Wolaver et al. 2008), creating unique conditions that remain underexplored in terms of microbial ecology (Souza et al. 2006). This study represents the first analysis of the microbial genome of the endangered Poza La Becerra spring, highlighting its untapped potential.

In the age of high-throughput sequencing, the cultivation of novel microorganisms continues to play an indispensable role in understanding microbial physiology, metabolism, and ecological roles (Lewis et al. 2021). This is especially true for extremotolerant microbes from unique ecosystems like CCB, where traditional culturing combined with genomic analysis can uncover microbial dark matter and valuable genetic pathways and phenotypic adaptations (Schultz et al. 2023).

*Streptomyces* species are well known for their ability to produce a diverse array of secondary metabolites, including antibiotics, anticancer agents, and immunosuppressants. However, genome mining approaches often prioritize genomes with high biosynthetic potential based on conventional criteria, which may inadvertently overlook microorganisms with unique biosynthetic capabilities (Gonzalez-Salazar *et al*. 2023). In particular, traditional bioinformatics tools tend to focus on genomes with a high number of biosynthetic gene clusters (BGCs) or those with well-characterized secondary metabolite pathways, neglecting potentially valuable candidates that may encode unconventional biosynthetic traits. Given the distinct adaptations of microorganisms to harsh conditions, the biosynthetic pathways of actinomycetes from Poza La Becerra could provide an untapped source of novel bioactive molecules with unique structural features.

Ribosomally synthesized and post-translationally modified peptides (RiPPs) have emerged as an exciting class of natural products due to their structural diversity, chemical stability, and broad-spectrum bioactivities, including antimicrobial, antiviral, and neuroactive properties. Unlike polyketides and nonribosomal peptides, RiPPs undergo extensive post-translational modifications that enhance their stability, specificity, and bioavailability, making them attractive candidates for drug development (Arnison *et al*. 2013; Han and Won 2024).

Recent computational models suggest that peptide sequences enriched in specific amino acids, particularly arginine, can exhibit enhanced penetration of blood-brain barrier (Gu *et al*. 2024).

Interestingly, extremophilic microorganisms, including those from high-stress environments such as CCB, are known to produce peptides with enriched arginine content, potentially conferring enhanced membrane permeability and neuroactive properties. This study reports the isolation, whole-genome sequencing, and genomic analysis of *Streptomyces* sp. PB17, a thermotolerant actinomycete from Poza La Becerra. Through genome mining, we identified 31 BGCs, including RiPP clusters with significant divergence from known pathways. We hypothesized that actinomycetes from extreme environments such as Poza La Becerra might encode RiPPs with unique structural features that could facilitate blood-brain barrier penetration, making them promising candidates for neurotherapeutic applications. To explore this potential, we employed predictive models to assess the likelihood of blood-brain barrier penetration and analyzed amino acid composition to identify sequence features associated with neuroactive properties. Our findings underscore the untapped potential of extremotolerant actinomycetes in the search for novel neurotherapeutics and highlight the importance of integrating genomic mining with predictive computational approaches in biodiscovery efforts.

## Materials and methods

### Study site, isolation and phenotypic characterization

Water and sediment samples were collected in Cuatro Ciénegas Basin from Pozas Becerras (26.87728º N, 102.13760º W) on march 8th, 2022. Access and permission to Natural Protected Area was given by SEMARNAT in document id.no. SGPA/DGVS/04225/21 issued on June 23th, 2021. On the day of sampling, temperature and pH were recorded at the site, yielding values of 30.5º and pH 6, respectively. Samples were transported at room temperature to the laboratory at CBTa No. 22 in Cuatro Ciénegas, Coahuila, where primary isolation was performed the same day. For selective isolation, ISP7 solid medium was used, prepared with the following composition ([per liter]): glycerol 15 g, L-tyrosine 0.5 g, L-asparagine 1 g, K_2_HPO_4_ (anhydrous basis) 0.5 g, MgSO_4_ · 7H_2_O 0.5 g, NaCl 0.5 g, FeSO_4_ · 7H2O 0.01 g, trace salts solution 1 mL, and agar 20 g. Fresh samples (150 µL) were plated onto the medium using sterile 3 mm glass beads for uniform spreading. Plates were incubated at 27° for 1–3 weeks. The PB17 isolate was selected based on distinct colony morphology and confirmed as an actinomycete through Gram staining. The isolate was re-streaked repeatedly until an axenic culture was obtained. The PB17 isolate was characterized by visual observation of macro and microscopic morphology in ISP2 medium ([per liter]: Yeast extract 4 g, Malt extract 10 g, Dextrose 4 g, Agar 20 g) using light microscopy with fresh and Gram stain properties. Morphological characterization of PB17 was conducted using ISP2 medium. Macro and microscopic observations were performed using light microscopy with fresh and Gram stained preparations. Growth temperature was assessed at four conditions (27°, 37°, 45° and 50°) to determine the optimal range. The isolate was stored in −20% glycerol at 20° for long term preservation and maintained at 4° for no longer than one month for subsequent analyses to find optimal ranges.

### DNA extraction and sequencing

Genomic DNA was prepared using a modified phenol/ chloroform method as described by Arocha-Garza *et al*. in 2017, DNA was quantified using the NanoDrop™2000 Fluorospectrometer (Thermo Scientific®NanoDrop; Thermo Fisher Scientific®, Wilmington, DE, USA) at a 260 nm absorbance, purity was measured using A260/280 absorbance ratio and integrity was assessed using 0.8% agarose gel. Whole genome was sequenced by Novogen (University of California at Davis, San Diego, Ca. USA) using Illumina NovaSeq 6000 × 150 at 100x depth.

### Assembly and quality assessment

Raw Sequencing data preprocessing was done partially in the Google colaboratory environment (Google 2024). As a first step, sequences were analyzed for quality using fastqc (Andrews 2016) and filtering and preprocessing was performed using fastp (Chen *et al*. 2018) applying several parameter combinations to remove adapters and miscalled bases, correct reads by overlap, base PHRED was established ≤ 20.

*De Novo* genome assembly was achieved using SPAdes v3.15.5 (Prjibelski *et al*. 2020) under default parameters. Quality assessment of assembly was done using Quast v5.2.0 (Gurevich *et al*. 2013) and CheckM v1.0.18 (Parks *et al*. 2015) and completeness was evaluated using Busco v5.7.1 at the Galaxy community server (Manni *et al*. 2021; Afgan *et al*. 2022) against the odb_bacteria10 gene family database under default parameters, using prodigal as gene predictor.

### Annotation and phylogeny

The assembly of PB17 was annotated with the Prokka software package (Seemann 2014) in the US departament knlowledgebase kbase (Arkin *et al*. 2018) as well as with BAKTA (Schwengers *et al*. 2021), and visualized using the Proksee Web server (Grant *et al*. 2023). Genome sequence data was uploaded to the Type (Strain) Genome Server (TYGS), a free bioinformatics platform available under https://tygs.dsmz.de, for a whole genome-based taxonomic analysis (Meier-Kolthoff and Göker 2019). Further-more, ANI calculations were performed using Kbase (Arkin *et al*. 2018). Two phylogenetic trees were constructed, first with the 20 validated core genes of the PB17 genome and related taxa recovered from NCBI, using the VBCG pipeline (Tian and Imanian 2023) and a second inferred by the GGDC web server (Meier-Kolthoff *et al*. 2022) available at http://ggdc.dsmz.de/ using the DSMZ phylogenomics pipeline (Meier-Kolthoff *et al*. 2014) adapted to single genes. A multiple sequence alignment was created with MUSCLE (Edgar 2004). Maximum likelihood (ML) and maximum parsimony (MP) trees were inferred from the alignment with RAxML (Stamatakis 2014) and TNT (Goloboff *et al*. 2008), respectively. For ML, rapid bootstrapping in conjunction with the autoMRE bootstopping criterion Pattengale *et al*. (2010) and subsequent search for the best tree was used; for MP, 1000 bootstrapping replicates were used in conjunction with tree-bisection-and-reconnection branch swapping and ten random sequence addition replicates. The sequences were checked for a compositional bias using the *χ*^2^ test as implemented in PAUP * (DL 2002). Trees obtained were visualized using iTOL (Letunic and Bork 2024).

### Biosynthetic gene clusters (BGCs), novelty scan and RiPPs analysis

Secondary metabolite Analysis Shell (antiSMASH) v7.1 (Blin *et al*. 2023) was applied to predict BGCs for secondary metabolite synthesis of strain PB17. The results obtained from the prediction were submitted to the BIG-FAM database (Kautsar *et al*. 2021) to determine the proximity of the query to gene cluster families and to calculate the biosynthetic novelty index (BiNI) using Eq. 1 as was established by González-Salazar *et al*. in 2023; let *d*_*i*_ be a sequence of proximity distances of the query to the database gene and *n*, the total number of predicted clusters, then:

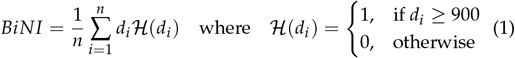

Additionally DeepBGC (Hannigan et al. 2019) was used to predict BGCs from the assembly annotations using deep learning. The tool’s default parameters and training data were utilized. The predicted features were added to the Proksee map to visualize in a genomic context and summarized using the R environment (R Core Team 2022; Wickham 2016).

#### Preliminar peptide analysis

The initial set of BGC predictions was narrowed to include only those related to RiPPs as these represent a relatively underexplored class of natural products with significant biotechnological and therapeutic potential. RiPPs are characterized by their structural diversity, smaller size, and modular biosynthesis, making them ideal candidates for bioengineering and optimization in drug discovery (Arnison *et al*. 2013). Their compact nature often enables them to traverse biological barriers, such as the blood-brain barrier, which is critical for addressing neurological diseases and infections. Additionally, RiPPs have been linked to a broad spectrum of bioactivities, including antimicrobial, antiviral, and anticancer properties, but remain underrepresented in genome mining studies compared to other classes like polyketides and nonribosomal peptides. The resulting subset was then processed using two distinct predictive models, both of which were designed to assess the likelihood of a peptide successfully penetrating the blood-brain barrier: SCMB3PP (Charoenkwan *et al*. 2022) and AUGUR (Gu *et al*. 2024). To consolidate predictions, results from both tools were visualized in a single scatter plot using R environment (R Core Team 2022; Wickham 2016; RStudio Team 2020). Subsequently, sequences were analyzed using GenomeSPOT (Barnum *et al*. 2024) to identify amino acid profile signatures associated with high expression in extremophiles. These signatures were then compared to amino acid frequencies reported in known Blood-Brain Barrier Penetrating Peptides (B3PPs). To examine those amino acid frequencies in RiPPs predicted by DeepBGC and AntiSMASH, a heatmap analysis (Hunter 2007; Van Rossum and Drake 2009; Wes McKinney 2010) and a Spearman’s correlation (Harrell Jr 2024; Wei and Simko 2024) were used.

## Results and discussion

### Morphological and phenotypic characterization of strain PB17

Strain PB17 showed morphological properties consistent with the *Streptomyces* genus. On the 7th day of growth, the isolate shows white aerial mycelium (RAL color standard [RAL] 3015) (für Gütesicherung und Kennzeichnung 1927), with a pink color (RAL 9018) in certain regions, which becomes more pronounced at day 9 with a reddish coloration in the sporulation zone (Figure 1). The complete morphological characterization of strain PB17 is provided in Supplementary Table S1 and Suplemmentary Figure S1. Optimal growth of *Streptomyces* sp. PB17 on solid media was observed at 37°, with robust growth persisting at 45° (Figure S2) suggesting potential thermotolerance.

**Figure 1.**
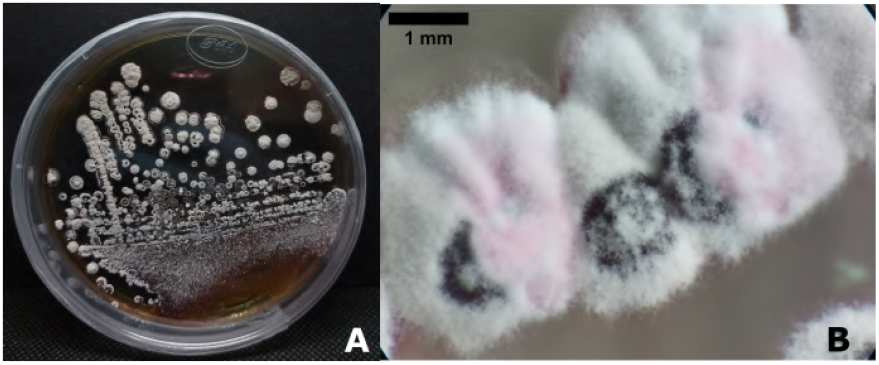
Macroscopic morphology of PB17 **A)** at 10 days of growth in ISP2 and **B)** magnification of colony at 7 days in ISP2.

### Genome analysis

Draft Genome of PB17 revealed the presence of a potential new *Streptomyces* species, as is shown by the phylogenomic analysis and the ANI calculation between PB17 and its related species (Supplementary Table S2), *Streptomyces tendae* JCM4610 (95.58%), *Streptomyces hyderabadensis* JCM17657 (93.98%) and *Streptomyces lividans* TK24 (93.72%). The most continuous contig corresponds to a length of 787,077 bp with a total G-C content of 72.19%. The genome was identified to be 99.7% complete with a 0.2% of missing families and 0.5% of duplications (Table 1). A map of genomic features for PB17 isolate can be seen in Figure S3. Genomic assemblies of high quality are indispensable for conducting in-depth analyses, such as the calculation of the BiNI (Gonzalez-Salazar *et al*. 2023) which are critical for exploring the unique capabilities of microorganisms like *Streptomyces* sp. PB17 as well as pangenomic studies. Species with the closest DNA–DNA relatedness based on digital DDH (dDDH) were *Streptomyces tendae* JCM 4610, *Streptomyces violaceoruber* CGMCC 4.1801, *Streptomyces rubrogriseus* NBRC 15455 and *Streptomyces anthocyanicus* JCM 5058 with values of 67.5, 67.6, 67.9 and 69.8% respectively (Supplementary Table S3). According to Komaki (2023), overclassification is a problem in the taxonomy of *Streptomyces*, however correlations between ANI, dDDH and MLSA evolutionary distance, along with BGCs clarification may help to avoid this problem, help with systematics and could be useful for novel bioactive metabolites.

**Table 1.** Genome assembly summary and annotation statistics for PB17 strain.

| Feature | PB17 |
| --- | --- |
| Total contigs | 70 |
| Largest contig | 787,077 |
| Total lenght | 8,714,003 |
| N50 | 301,447 |
| L50 | 11 |
| L90 | 31 |
| GC (%) | 72.19 |
| Mismatches<sup>a</sup> | 0 |
| tRNA | 70 |
| rRNA | 4 |
| CDS | 8091 |
| CHECKM Completeness | 100% |
| Total BUSCO's group searched<sup>b</sup> | 1579 |
| Complete BUSCO's | 1575 |
| single copy | 1567 |
| duplicated copy | 8 |
| Fragmented BUSCO's | 2 |
| Missing BUSCO's | 2 |

<sup>a</sup> as defined by average number of uncalled bases (N's) in 100kbp.<sup>b</sup> against odb\_bacteria10

The phylogenetic analyses of *Streptomyces* sp. PB17 were conducted using two complementary approaches: the Type (Strain) Genome Server (TYGS) and the Validated Bacterial Core Genes (VBCG) pipeline. For the inferred tree with TYGS, the input nucleotide matrix comprised 19 operational taxonomic units and 1525 characters, 50 of which were variable and 20 of which were parsimony-informative. The base-frequency check indicated no compositional bias (p = 1.00, *α* = 0.05). ML analysis under the GTR+GAMMA model yielded a highest log likelihood of −2516.02, whereas the estimated alpha parameter was 0.02. The ML bootstrapping did not converge, hence 1000 replicates were conducted; the average support was 50.75%. MP analysis yielded a best score of 62 (consistency index 0.89, retention index 0.86) and 60 best trees. The MP bootstrapping average support was 31.56% Figure 2

**Figure 2.**
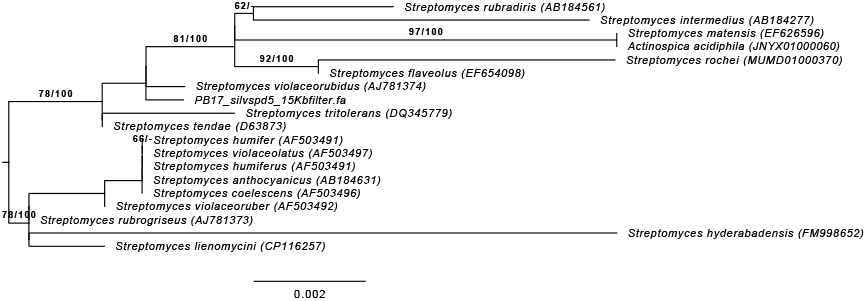
ML tree inferred under the GTR+GAMMA model and rooted by midpoint-rooting The branches are scaled in terms of the expected number of substitutions per site. The numbers above the branches are support values when larger than 60% from ML (left) and MP (right) bootstrapping.

On the other hand, for consider not only high presence and single-copy ratios but also the genes’ phylogenetic fidelity—a measure of the consistency of phylogenies within the 20 core gene set was evaluated in the tree inferred by VBCG (Tian and Imanian 2023), as is shown in Figure 3. Upon visual inspection, the trees exhibit certain topological differences; however, both pipelines consistently placed the isolate within the genus *Streptomyces*.

**Figure 3.**
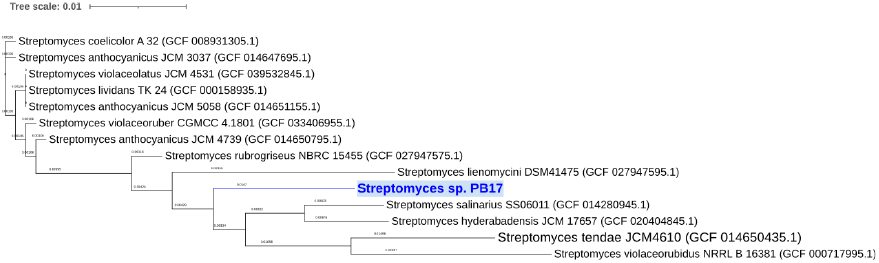
VBCG tree of the 20 concatenated core genes of each genome.

The congruence between the ANI results and the finer resolution of the VBCG tree strongly supports the potential phylogenetic novelty of *Streptomyces* sp. PB17. Although the TYGS phylogeny offers a broader perspective of genomic similarity within the genus, the higher resolution of the VBCG approach provides more reliable evidence for species-level delineation.

### Biosynthetic Novelty

The result of the gene cluster prediction by applying antiSMASH showed that strain PB17 contained 31 Biosynthetic clusters (2 RiPPs), among them 10 clusters exhibited low similarity to known characterized pathways (Table 2), representing space for new metabolites, as determined by BiNI analysis. Although the BiNI value (374.48) of the PB17 genome indicates moderate divergence from well-characterized pathways, as was stated by González-Salazar *et al*., the biosynthetic potential of this novel *Streptomyces* species from the CCB, underscores its capacity to contribute unique chemical diversity, particularly as a thermoresistant strain adapted to an extreme environment.

**Table 2.** Biosynthetic Gene Clusters (BGC) distribution of potential novelty found by AntiSMASH (v 7.1) predictions.

| Type<sup>a</sup> | Product | Length (Kb) | Similar to: | % Similarity |
| --- | --- | --- | --- | --- |
| Polyketide: T3PKS | Butyrolactone | 49.59 | Germicidin | 100% |
| Other: NI | Siderophore | 29.79 | desferrioxamin B/ des-ferrioxamine E | 100% |
| RiPP:RRE-containing | Terpene | 21.24 | naphthomycin A | 9% |
| NRPS-like |  | 41.42 | Aurantimycin A | 20% |
| Other/ Uncategorized | Unknown | 40.89 | ND | ND |
| NRP | Metallophore | 58.44 | coelichelin | 100% |
| NRPS |  | 26.85 | Polyoxypeptin | 16% |
| Polyketide: T1PKS |  | 19.34 | polyoxypeptin | 18% |
| Other NI | Siderophore | 31.21 | Enduracidin | 10% |
| Unknown | hydrogen-cyanide | 12.98 | Aborycin | 21% |

<sup>a</sup> BGC type in this table were those whose $d_i \geq 900$ or $\mathcal{H}(d_i) = 1$ (Equation 1) and that possibly express novelty products

A notable increase in BGCs number (152) was detected using the deep learning prediction model (DeepBGC), the tool predicted, among other classes, 18 PKS, 11 RiPPs and 102 new biosynthetic gene clusters, with predominantly antibacterial activity observed throughout the draft genome (Supplementary Figures S4 y S5).

### RiPPs

*Streptomyces sp*. PB17 exhibited notable scores for B3PP across the predictive tools used. From the initial pool of 13 RiPPs, as predicted by DeepBGC and AntiSMASH, three were determined to have a high probability of penetrating the blood-brain barrier based on their B3PP scores generated by AUGUR and SCM3BPP (Figure 4).

**Figure 4.**
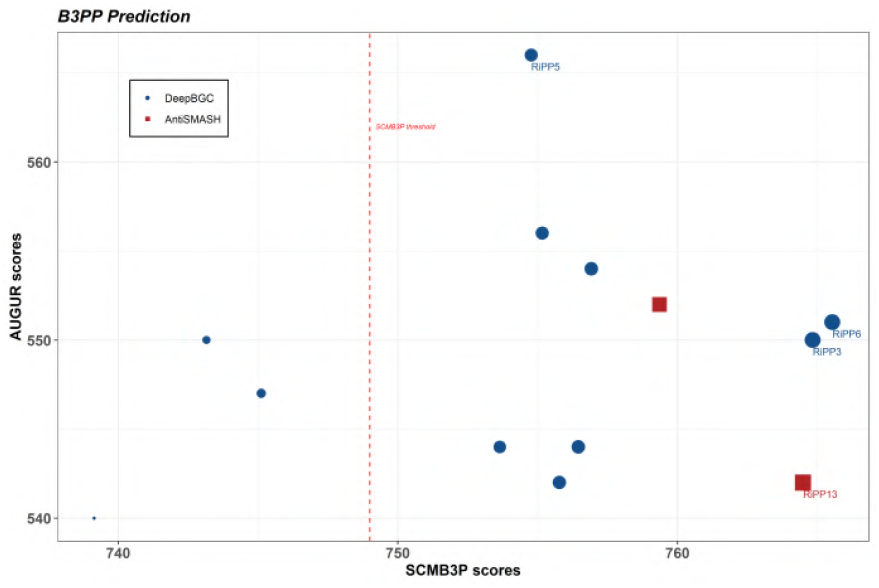
Scatter plot of predictive scores by tool (blue points corresponding to deepBGC and red square Antismash) for Blood-Brain barrier penetrating peptides according to AUGUR (y axis) and SCM3BPP (x axis). Red line represents the thresh-old limit for positive prediction in SCM3BPP ((Charoenkwan *et al*. 2022). According to AUGUR all the graph area represent positive predictions

Among PB17 predicted RiPPs, RiPP6 exhibited the highest score in SCMB3PP (765.54) and RiPP5 shows the highest score in AUGUR (560). Bio_PFAM examination revealed that RiPP6 is associated with PF02668, a domain commonly found in TauD enzymes involved in taurine catabolism and is encountered in 1104 Streptomycetaceae species while RiPP5 is with PF08242, PF08241 Methyltransferase domains and PF01488 a Shikimate/ quinate 5-dehydrogenase (Mistry *et al*. 2021). RiPP3 (predicted by Deep-BGC) and RiPP13 (predicted by AntiSMASH) also showed high B3PP scores in both tools (Figure 4). Notably, both DeepBGC and AntiSMASH independently predicted RiPPs at the same genomic node as is shown in Table 3, suggesting high confidence in the predicted RiPP. While these tools indicate a strong potential for novel bioproducts, blood-brain barrier penetration, the precise biological function of these putative BGCs remains uncertain without custom-trained models, data augmentation and experimental testing.

**Table 3.** Characteristics of RiPPs with B3PP consensus prediction and highest scores among tools.

| B3PP RiPP | Prediction tool | Length (pb) | Proteins <sup>a</sup> | Domains | Location |
| --- | --- | --- | --- | --- | --- |
| RiPP3 | DeepBGC | 2,970 | 2 | 10 | NODE6 |
| RiPP5 | DeepBGC | 4,756 | 6 | 24 | NODE7 |
| RiPP6 | DeepBGC | 4,288 | 4 | 10 | NODE8 |
| RiPP13 | AntiSMASH | 10,216 | 5 | ND<sup>b</sup> | NODE6 |

<sup>a</sup> Putative proteins or genes in prediction
<sup>b</sup> This information is absent from the AntiSMASH output

These RiPPs predictions, particularly RiPP6, are unprecedent genomic evidence for potential blood-brain barrier penetrating molecules. At sequence evaluation level, since peptides produced by extremophiles display an enriched frequency of certain aminoacids (Panja *et al*. 2020), aminoacidic profiling was analyzed from PB17, revealing that the pool of predicted RiPPs are rich in arginine (Arg), followed by alanine (Ala), glycine (Gly), and proline (Pro). The highest abundance of Arg was observed in RiPP9 (0.27), followed by RiPP1 (0.21), with the remaining RiPPs exhibiting moderate Arg values ranging from 0.07 to 0.2. Alanine ranged from 0.099 to 0.17, glycine from 0.076 to 0.16, and proline from 0.048 to 0.17. In contrast, asparagine (Asn), tyrosine (Tyr), and methionine (Met) were present at low abundance. Among the RiPPs with the highest B3PP scores, a moderate abundance of Arg (not exceeding 0.2) was observed, with alanine frequencies ranging from 0.13 to 0.17, followed by Gly (0.07-0.15). Very low frequencies of tryptophan (Trp) (0.005-0.01), Asn (0.003-0.012), and Met (0.004-0.017) were also noted. A heatmap of amino acid frequencies is available in Supplementary Figure S6. Moreover, analysis of the calculated valency (Zc) using GenomeSPOT revealed that the RiPPs predicted with the highest B3PP scores exhibit negative or slightly positive (near-neutral) charges, with values ranging from -0.1 to 0.04.

To date, no studies have explored RiPPs of microorganisms from extreme sites as potential sources of B3PPs, however, genomic signature analyses have revealed that extremophilic bacteria, thriving in extreme conditions of temperature, pH, and salinity, exhibit elevated expression of specific amino acids, including Arg, Gly, Ala, Pro and aspartic acid (Asp) (Arias *et al*. 2023; Loukas 2024) and notably, Arg and tyrosine (Tyr) are overrepresented in known B3PPs, while Gly, Pro, Lysine (Lys), and leucine (Leu) are moderately enriched compared to other amino acids (Gu et al. 2024). For instance, some of the chemical structure of these aminoacids, such as the positive charge of Arg and its ability to form hydrogen bonds can contribute to its high prevalence in B3PPs. The blood-brain barrier is negatively charged, limiting the permeability of negatively charged molecules while favoring positively charged ones (Gu et al. 2024). This suggests that Arg, with its positive charge, plays a crucial role in enabling B3PPs to traverse the blood-brain barrier. Particularly, the observed high frequency of Arg in PB17 was also noticed in at least two other actinomycetal isolates from CCB (data not shown). This adaptation could provide stability to proteins under harsh conditions or play a role in osmo or thermoregulation (Siddiqui *et al*. 2006; Khan and Patra 2018; Barnum et al. 2024), additionally, Arg has been implicated in antibiotic production in microorganisms (Zhang et al. 2020; Eloïse et al. 2024). In this sense, the antibiotic tirandamycin B, produced by *Streptomyces composti* sp. nov., has been found to inhibit *β*-secretase 1, a key player enzyme in Alzheimer’s disease, suggesting potential neuroprotective benefits (Duangupama et al. 2023).

Spearman correlation analysis between amino acid frequencies across the set of PB17 predicted RiPPs and both B3PP predictive tools was performed (Supplementary Figure S7), revealing that AUGUR predictions showed a strong positive correlation with Ala frequency (Spearman *ρ* = 0.7614, *p* ≤ 0.005) and a negative correlation with Trp frequency (Spearman *ρ* = −0.56, *p* ≤ 0.04). SCMB3PP predictions were more strongly correlated with a greater number of amino acids: negatively with Arg, Ser, Cys, and Trp frequencies, and positively with Val, Asp, Gly, Phe, Gln, Glu, and Leu frequencies (Figure S7). Both tools concurred in the negative correlation with Trp frequency. No direct correlation was observed between a higher arginine abundance and higher B3PP scores. For instance, RiPP 9 from *Streptomyces sp*. PB17, which exhibited the highest Arg frequency among the predicted RiPPs, was categorized as a non-B3PP by SCMB3PP. This discrepancy could be explained by the potential destabilizing effect of excessive Arg, as the guanidinium group in this aminoacid can disrupt peptide structure (Ren 2023). This suggests that, for the specific RiPPs in this strain, a non-excessive amount of Arg may favor positive B3PP scores. Given the nature of the prediction algorithms, with SCMB3PP being a scoring card method-based predictor that considers dipeptide propensities but with limited attention to the information imbalance between B3PPs and non-B3PPs, and without using data augmentation methods (Charoenkwan et al. 2022; Gu et al. 2024), these results should be interpreted with caution. However, both tools concurred in the prediction of RiPPs with the potential to cross the blood-brain barrier.

While this study provides compelling computational evidence supporting the potential of *Streptomyces* sp. PB17 as a source of neuroactive RiPPs, our findings are based on predictive analyses and require experimental validation. Functional characterization, biochemical assays, *in vitro* and *in vivo* studies are essential next steps to confirm the biological activity and blood-brain barrier penetration capacity of the identified peptides. However, these experiments require significant time, resources, and specialized methodologies, which pose challenges in advancing this work to the next stage of the biodiscovery pipeline. Despite these limitations, our study establishes a strong foundation for guiding future experimental efforts and broadening the scope of natural product discovery from extremophilic microorganisms. Further research is needed to establish a definitive cause-and-effect relationship. Collectively, these findings open new possibilities for exploring microorganisms from extreme environments and their secondary metabolites in the context of neurotherapeutic application.

## Conclusion

To date, there has been no reported attempt to predict potentially neuroactive RiPPs from microbial genomes, making this study the first to explore this avenue. Our findings underscore the untapped potential of extremophilic *Streptomyce*s strains from underexplored environments, such as Poza La Becerra in the Cuatro Ciénegas Basin, in the search for novel neurotherapeutics. The genome mining of *Streptomyces* sp. PB17 revealed RiPP biosynthetic gene clusters with significant divergence from known pathways, suggesting the presence of structurally unique bioactive molecules. Furthermore, predictive analyses identified several RiPPs with high probability scores for blood-brain barrier penetration, reinforcing their potential relevance for neurotherapeutic applications.

Beyond expanding the known biosynthetic repertoire of *Streptomyces*, this study challenges the prevailing biases in genome mining. By leveraging predictive models, deep-learning tools, and unconventional screening strategies, we demonstrate that valuable biosynthetic potential can reside even in genomes that may otherwise be overlooked due to moderate BiNI values or lower counts of classical BGCs.

Ultimately, our findings highlight the importance of reconsidering microbial genomes that appear to be well-characterized or of low biosynthetic novelty under standard screening parameters. By integrating computational predictions with genomic analysis, this approach paves the way for a more inclusive and systematic exploration of microbial biodiversity, with direct implications for pharmaceutical discovery and neurotherapeutic development.

## Data availability

This Whole Genome Shotgun project has been deposited at DDBJ/ENA/GenBank under the accession JBJPFQ000000000 associated with the BioProject ID PRJNA1193220. The version described in this paper is version JBJPFQ010000000. Supplementary material can be found at https://doi.org/10.6084/m9.figshare.27950112.v1.

## Acknowledgment

The authors would like to thank Alejandra Arreola Triana for her valuable assistance in revising the manuscript and providing insightful suggestions. The authors also thank Cynthia D. Correa-Oviedo for her exceptional technical support in the laboratory, which was instrumental in the successful completion of this work. The authors used ChatGPT and Gemini to ensure proper grammar and language use of this manuscript. After using these tools, the authors checked the work to ensure accuracy and are fully responsible for the content of the publication.

